# Evidence for a role of 5-HT-glutamate co-releasing neurons in acute stress mechanisms

**DOI:** 10.1101/2023.11.25.568634

**Authors:** L. Sophie Gullino, Cara Fuller, Poppy Dunn, Helen M. Collins, Salah El Mestikawy, Trevor Sharp

## Abstract

The majority of midbrain 5-hydroxytryptamine (5-HT) neurons express the vesicular glutamate transporter 3 (VGLUT3) and co-release 5-HT and glutamate, but the function of this co-release is unclear. Given the strong links between 5-HT and uncontrollable stress, we used a combination of c-Fos immunocytochemistry and conditional gene knock out in mice to test the hypothesis that glutamate co-releasing 5-HT neurons would be activated by stress and involved in stress coping.

Acute, uncontrollable swim stress increased c-Fos immunoreactivity in neurons co-expressing VGLUT3 and the 5-HT marker tryptophan hydroxylase 2 (TPH2) in the dorsal raphe nucleus (DRN). This effect was localised in the ventral DRN subregion and prevented by the antidepressant fluoxetine. In contrast, a more controllable stressor, acute social defeat, had no effect on c-Fos immunoreactivity in VGLUT3-TPH2 co-expressing neurons in the DRN.

To test whether activation of glutamate co-releasing 5-HT neurons was causally linked to stress coping, mice with a specific deletion of VGLUT3 in 5-HT neurons were exposed to acute swim stress. Compared to wildtype controls, the mutant mice showed increased climbing behaviour, a measure of active coping. Wildtype mice also showed increased climbing when administered fluoxetine, revealing an interesting parallel between the behavioural effects of genetic loss of VGLUT3 in 5-HT neurons and 5-HT reuptake inhibition.

We conclude that 5-HT-glutamate co-releasing neurons are recruited by exposure to uncontrollable stress. Furthermore, natural variation in the balance of 5-HT and glutamate released at the 5-HT synapse may impact on stress susceptibility.

## Introduction

Serotonin (5-hydroxytryptamine; 5-HT) is a key neuromodulator of emotional processing, stress sensitivity and coping behaviour^1,2^. 5-HT neurons in the midbrain dorsal raphe nucleus (DRN), the principal source of 5-HT innervation to the forebrain, are activated by acute inescapable stressors such as forced swim, restraint and foot-shock, as evident through increased expression of the activity-dependent immediate-early gene *c-fos* in 5-HT neurons^3–8^. Although other forms of stress also activate 5-HT neurons^9,10^, evidence suggests that stressors allowing for the least control (i.e. inescapable stressors) are associated with greater 5-HT neuron activation^9,11,12^.

Recently, it has become clear that 5-HT neurons are capable of releasing not only 5-HT but also glutamate. Electrophysiological evidence for 5-HT-glutamate co-release in cultured 5-HT neurons^13^ was followed by the discovery of the expression of the type 3 vesicular glutamate transporter (VGLUT3) in the majority (50-80 %) of 5-HT neurons^14–16^. More recently, electrophysiological studies have demonstrated that optogenetic activation of 5-HT neurons elicits both 5-HT and glutamate-mediated synaptic responses in different forebrain regions^17–19^.

Currently the functional role of 5-HT-glutamate co-release is unclear although links to anxiety-like behaviour and reward processing have been proposed based on studies of the phenotype of VGLUT3 knockout mice^19–21^ and the behavioural effects of optogenetic activation of 5-HT neurons^19,20^. Interestingly, in a recent chemogenetic study, activation of 5-HT neurons projecting to the prefrontal cortex from the ventral region of the DRN, an area rich in 5-HT-glutamate co-releasing neurons, increased active coping (i.e. reduced immobility) in mice exposed to swim stress^22^. The latter finding suggests that glutamate co-releasing 5-HT neurons are activated by uncontrollable stressors, such as swim stress, and may be involved in stress coping behaviour. This result^22^ also emphasises the functional heterogeneity within DRN subregions that has been detected in previous studies^8,23,24^.

Here we used c-Fos immunohistochemistry to test the prediction that 5-HT-glutamate co-releasing neurons in the DRN (particularly the ventral region) would be activated by an uncontrollable stressor, specifically swim stress. Effects were compared with a more controllable stressor, acute social defeat. Finally, behavioural experiments using a novel transgenic mouse with VGLUT3 knockout targeted to 5-HT neurons (VGLUT3 cKO^5-HT^ mice^25^) examined the causal link between changes in activity of 5-HT-glutamate co-releasing neurons and stress coping behaviour.

**Figure 1.**
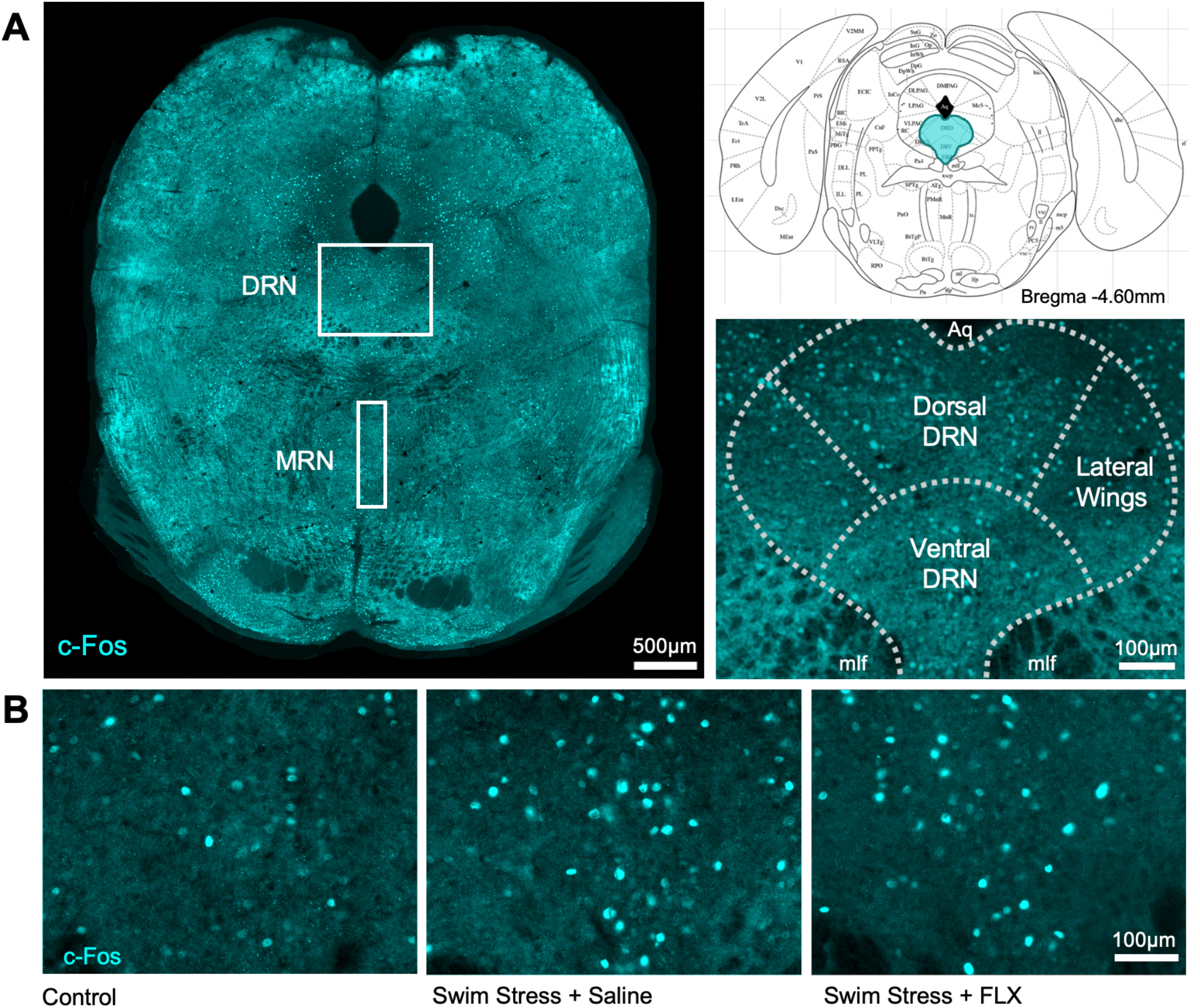
C-Fos immunoreactivity in mouse midbrain following acute swim stress. (A) C-Fos immunoreactivity in a midbrain section at the level of the DRN and MRN (left) according to the stereotaxic atlas (top right) of ^26^. Higher magnification images of DRN subregions (bottom right). (B) High magnification images of c-Fos immunoreactivity in the ventral DRN of control mice, and mice administered a single injection of either saline or fluoxetine (FLX) and exposed to swim stress. Abbreviations: dorsal raphe nucleus (DRN), median raphe nucleus (MRN), aqueduct (Aq), medial longitudinal fasciculus (mlf).

## Results and Discussion

### Swim stress evoked c-Fos expression in the DRN

Exposure of mice to acute swim stress increased the number of c-Fos immunoreactive neurons in the DRN and median raphe nucleus (MRN) (effect of treatment: F_(2,17)_=5.503, p=0.014; effect of region: F_(1,15)_=17.160, p<0.001; region x treatment interaction: F_(2,15)_=0.272, p=0.766; ***Fig. 2A***). Post-hoc analysis revealed that this effect was statistically significant in the DRN of swim-stressed mice compared to non-stressed controls (p=0.017; ***Fig. 2A***). Conversely, the number of c-Fos immunoreactive cells in the MRN was not significantly different across conditions (F_(2,15)_=2.065, p=0.161; ***Fig. 2A***). These data are in accord with previous studies reporting that swim stress increased c-Fos immunoreactivity in DRN of rats^8,23^.

**Figure 2.**
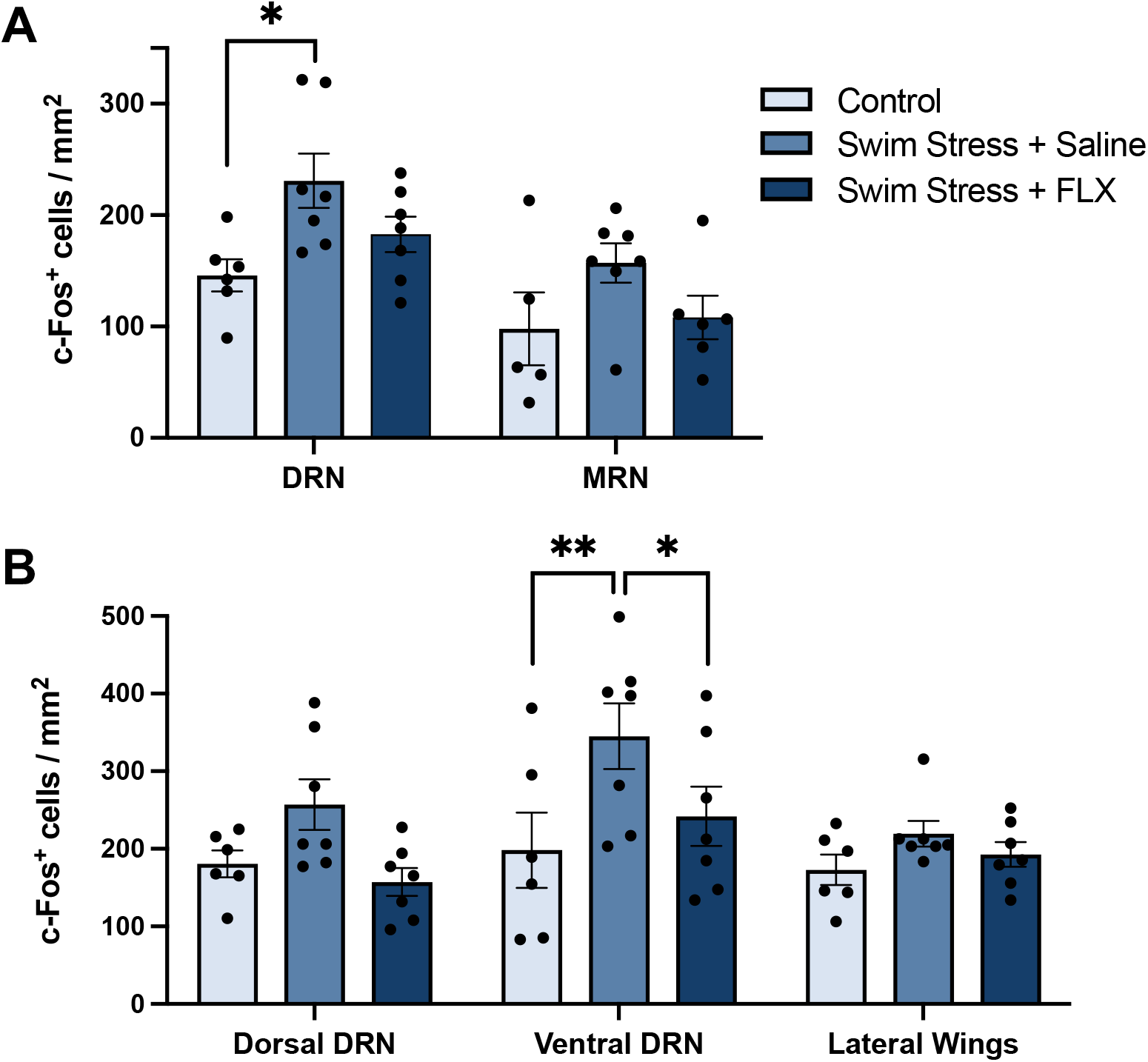
Effect of acute swim stress, with or without fluoxetine, on c-Fos expression in midbrain subregions. (A) C-Fos immunoreactive neurons in the DRN and MRN. (B) C-Fos immunoreactive neurons in DRN subregions. Columns are mean ± SEM values with individual values indicated by closed circles. ** p<0.01 *p<0.05. Groups were control (n=6), saline + swim stress (n=7) and 10 mg/kg fluoxetine + swim stress (n=7). Abbreviations as in Fig. 1.

Further examination of the DRN at the subregional level (***Fig. 2B***) revealed a statistically significant effect of both region (F_(2,34)_=5.884, p=0.006) and treatment (F_(2,17)_=5.721, p=0.013). Although the region x treatment interaction was not statistically significant (F_(4,34)_=1.512, p=0.221), likely due to the small sample size, post-hoc testing was deemed justified based on previous evidence and our a priori hypothesis of a preferential involvement of ventral DRN neurons in stress coping (see Introduction). Post-hoc analysis showed a statistically significant increase in c-Fos immunoreactive neurons in the ventral DRN of swim-stressed mice compared to non-stressed controls (p=0.002; ***Fig. 2B***), but non-significant effects in the dorsal DRN (p=0.181) and lateral wings (p=0.520). Pre-treatment with the selective serotonin reuptake inhibitor (SSRI) fluoxetine (10 mg/kg i.p.) prevented stress-induced c-Fos expression in the ventral DRN (post-hoc p=0.028; ***Fig. 2B***). Additionally, during swim stress fluoxetine-treated mice spent more time climbing, a measure of active coping (Mann-Whitney U=6, p=0.016; ***Suppl. Fig. 1***; see later for further discussion).

### Swim stress increased c-Fos expression in DRN neurons co-expressing TPH2 and VGLUT3

Next, we investigated whether swim stress increased c-Fos immunoreactivity specifically in 5-HT-glutamate co-releasing neurons, using the same sections examined for c-Fos alone. Previous studies have revealed that VGLUT3-expressing neurons in the midbrain raphe nuclei comprise two sub-populations, one colocalizing 5-HT and the other lacking 5-HT^16,27^. Here, the 5-HT-specific marker tryptophan hydroxylase 2 (TPH2) was used to distinguish these two populations (***Fig. 3A***). In agreement with these earlier studies, somatic VGLUT3 expression was particularly evident in TPH2 immunoreactive neurons located in the ventral DRN; thus, 67.9 ± 3.04 % of TPH2 immunoreactive neurons co-expressed VGLUT3 (***Suppl. Fig. 2***). In comparison, only sparse VGLUT3 expression was observed in TPH2 immunoreactive neurons in the dorsal DRN and lateral wings. Neurons with colocalised VGLUT3 and TPH2 were evident in the MRN although these neurons were less abundant than the DRN; thus, 34.9 ± 2.9 % of TPH2 immunoreactive neurons also expressed VGLUT3.

**Figure 3.**
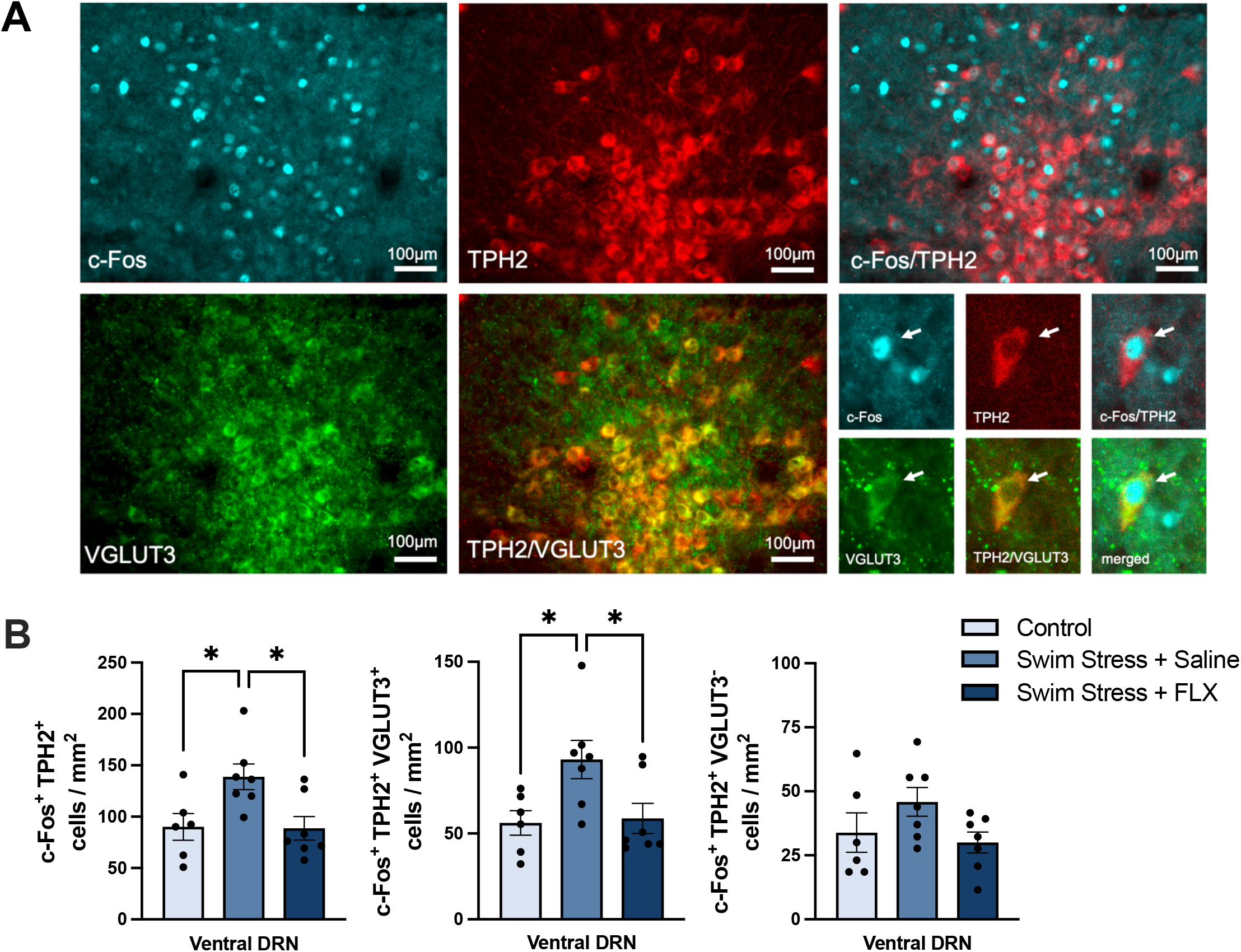
Effect of swim stress, with or without fluoxetine, on c-Fos expression in DRN neurons co-expressing TPH2 and VGLUT3 in the ventral DRN. (A) Representative image of c-Fos/TPH2/VGLUT3 triple-labelled neurons in the ventral DRN (AP= −4.6mm). (B) Effect of swim stress on number of c-Fos/TPH2 double-labelled neurons (left), c-Fos/TPH2/VGLUT3 triple-labelled neurons (middle), and c-Fos/TPH2 double-labelled neurons but VGLUT3 immunonegative (right). Columns represent the mean ± SEM values, with individual values indicated by closed circles. * p<0.05. Groups were control (n=6), saline + swim stress (n=7) and 10 mg/kg fluoxetine + swim stress (n=7). Abbreviations as in Fig. 1.

Importantly, swim stress increased the number of c-Fos/TPH2/VGLUT3 triple-labelled neurons in the ventral DRN compared to non-stressed controls (F_(2,17)_= 4.896, p=0.021, post-hoc p=0.036; ***Fig. 3B***). This effect of swim stress amounted to an increase in c-Fos in 32.3 ± 7 % of TPH2/VGLUT3 immunoreactive neurons in the ventral DRN. Furthermore, compared to saline controls pre-treatment with fluoxetine prevented the stress-induced increase in c-Fos immunoreactivity in TPH2/VGLUT3 co-expressing neurons (post-hoc p=0.042; ***Fig. 3B***).

Swim stress also significantly increased the number of c-Fos/TPH2 double-labelled neurons in the ventral DRN (F_(2,17)_=5.535, p=0.014, post-hoc p=0.034) compared to non-stressed controls (26.1 ± 2.8 % of TPH2 immunoreactive neurons), and this effect was also reduced by fluoxetine (F_(2,17)_=5.535, p=0.014, post-hoc p=0.023; ***Fig. 3B***). TPH2 immunoreactive neurons that were immunonegative for VGLUT3 did not show increased c-Fos expression in response to swim stress (F_(2,17)_= 2.115, p=0.151; ***Fig. 3B***). The number of TPH2 immunoreactive neurons did not differ between groups (***Suppl. Fig. 3A***).

To our knowledge this is the first report of evidence that 5-HT-glutamate co-releasing neurons in the ventral DRN are activated by exposure to a stressor, specifically acute swim stress. The inhibitory effect of fluoxetine on this stress-evoked response is in line with electrophysiological evidence that acute SSRI administration inhibits the firing of DRN 5-HT neurons through a 5-HT_1A_ autoreceptor mediated hyperpolarisation^28–30^.

### Social defeat did not evoke c-Fos expression in DRN neurons co-expressing TPH2 and VGLUT3

Previous c-Fos studies report that 5-HT neurons in the DRN are more sensitive to uncontrollable versus controllable stressors^9,11,12,31^. Acute swim stress is a well-established inescapable stressor, whereas social defeat is an example of a more controllable stressor. Thus, socially-defeated animals adopt a variety of active coping strategies (e.g. flight, corner location, upright submissive postures) to minimise interactions with the opponent^32^.

We utilised the social defeat model to investigate the sensitivity of VGLUT3-expressing 5-HT neurons to a more controllable stressor. Here, naive intruder mice were exposed to a single episode of social defeat in the home cage of a larger territorially-dominant resident. Socially defeated mice were separated from the resident after a single defeat episode that was typically limited to less than a 1 min to avoid the stressor from becoming inescapable. The average latency for the resident to attack was 5.1 ± 1.7 s and the average number of attacks per encounter was 14.9 ± 2.8, i.e. an attack every 3 s, involving a combination of biting, kicking, and wrestling, prior to a clear pin down (social defeat). During the encounter intruder mice spent most of the time moving (90 ± 3.1 %) and actively avoiding the resident (distance travelled 3.4 ± 0.8 m).

Region-specific analysis showed that acute social defeat had no effect on the number of c-Fos immunoreactive neurons in the ventral DRN compared to non-stressed controls, and other DRN subregions were similarly unaffected (effect of region: F_(1.815,24.50)_=0.822, p=0.441, effect of treatment: F_(1,14)_=0.064, p=0.804, treatment x region interaction F_(2, 27)_= 1.123, p=0.340; ***Fig. 4A***). Moreover, the number of c-Fos/TPH2 double-labelled neurons in the ventral DRN was not different across groups (t_(13)_=1.158, p=0.403; ***Fig. 4B***). Importantly, and in contrast to swim stress, acute social defeat did not alter the number of c-Fos/TPH2/VGLUT3 triple-labelled neurons in the ventral DRN compared to non-stressed controls (t_(13)_=0.732, p=0.167; ***Fig. 4B***).

**Figure 4.**
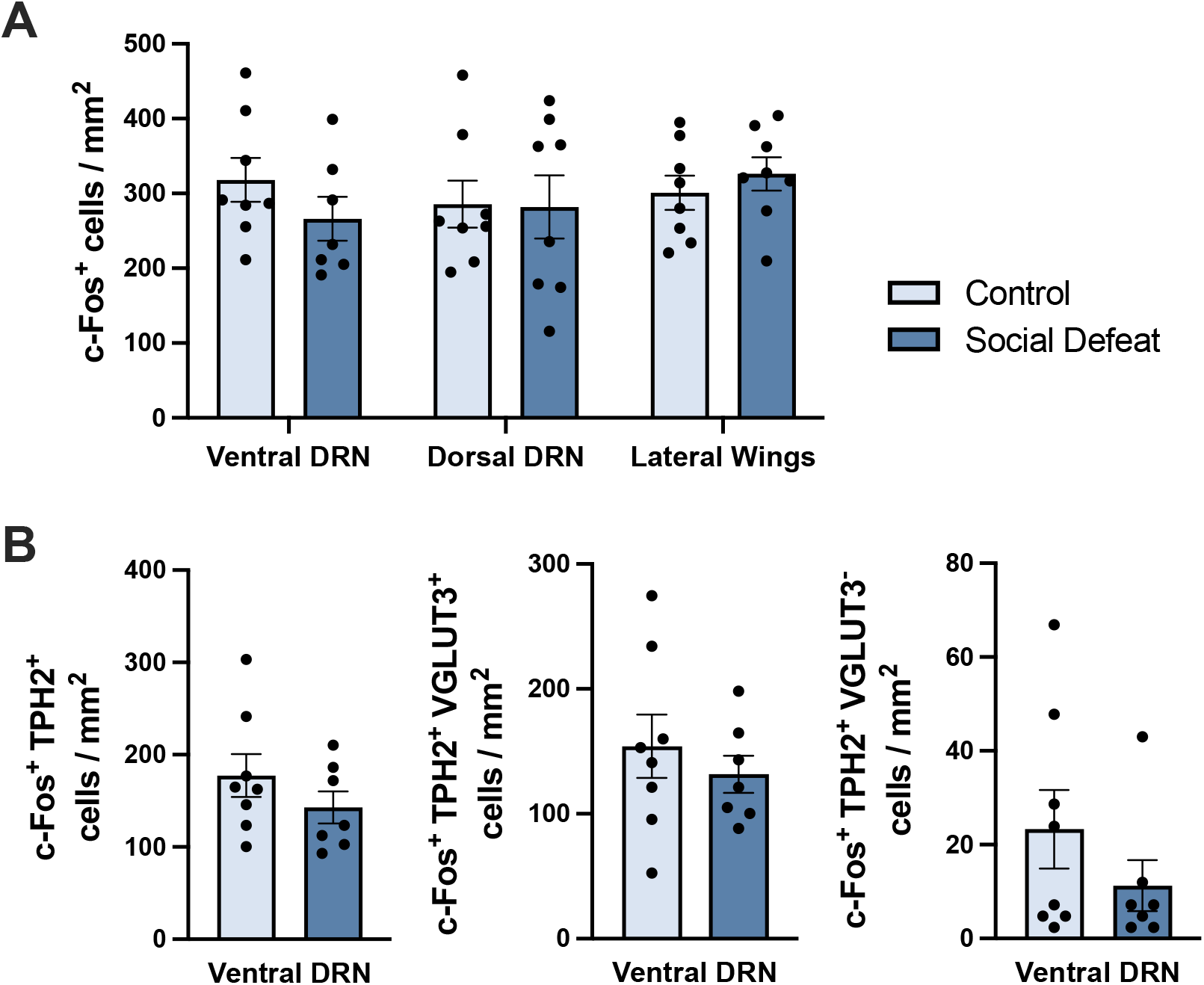
Effect of acute social defeat on c-Fos immunoreactive neurons in DRN, including neurons co-labelled with TPH2 and VGLUT3. (A) C-Fos immunoreactive neurons in DRN subregions. (B) C-Fos/TPH2 double-labelled neurons (left), c-Fos/TPH2/VGLUT3 triple-labelled neurons (middle), and c-Fos/TPH2 double-labelled neurons immunonegative for VGLUT3 (right) in the ventral DRN. Columns represent mean ± SEM values, with individual values indicated by closed circles. Groups were non-stressed controls (n=8) and social defeat (n=7). Abbreviations as in Fig. 1.

Social defeat also had no effect on c-Fos expression in TPH2 neurons which were VGLUT3 immunonegative (t_(13)_=1.167, p=0.264; ***Fig. 4B***), and the number of TPH2 immunoreactive neurons in the ventral DRN was also unchanged (***Suppl. Fig. 3B***).

The lack of effect of social defeat on c-Fos expression in the DRN is in line with previous studies exposing rodents to a single short (∼3 min) period of social defeat^33,34^. Although some studies report that acute social defeat increased c-Fos expression in DRN neurons^35,36^, these findings were obtained from animals exposed to the resident over a long period (∼10 min) such that the stressor likely becomes inescapable^33^

Thus, the current data suggest that 5-HT-glutamate co-releasing neurons are preferentially activated by an uncontrollable versus controllable stressor. These data agree with previous c-Fos studies reporting that 5-HT neurons are more sensitive to uncontrollable versus controllable footshock^9,31^ but extend the findings to 5-HT-glutamate co-releasing neurons. Based on previous experiments involving localised muscimol injections it was concluded that controllable stressors have less impact on DRN 5-HT neurons due to the inhibitory influence of the medial prefrontal cortex^11^. Thus, the greater effect of swim stress versus social defeat on VGLUT3-expressing 5-HT neurons could be explained by the same mechanism.

It could be argued that the lack of effect of social defeat on DRN neurons is due to the strength of the stressor being insufficient. However, social defeat increased c-Fos expression in the periaqueductal gray (PAG). Thus, in socially defeated mice, c-Fos expression increased in the dorsal PAG compared to non-stressed controls (effect of region: F_(1,14)_=181.4, p<0.0001, effect of treatment: F_(1,14)_=10.20, p=0.007, region x treatment interaction: F_(1,14)_=8.358, p=0.012, post-hoc p=0.001; ***Fig 5A***), and there was a trend effect in the ventrolateral region (p=0.081). In comparison, swim stress also increased c-Fos expression in the dorsal and ventrolateral PAG (effect of region: F_(1,10)_=60.77, p<0.0001, effect of treatment: F_(1,10)_=58.78, p<0.0001, region x treatment interaction: F_(1,10)_=5.597, p=0.04, post-hoc p=0.001 and p<0.0001; ***Fig 5B***). PAG sub-regions are well known to be activated by stress^37^ and involved in stress coping^38,39^.

**Figure 5.**
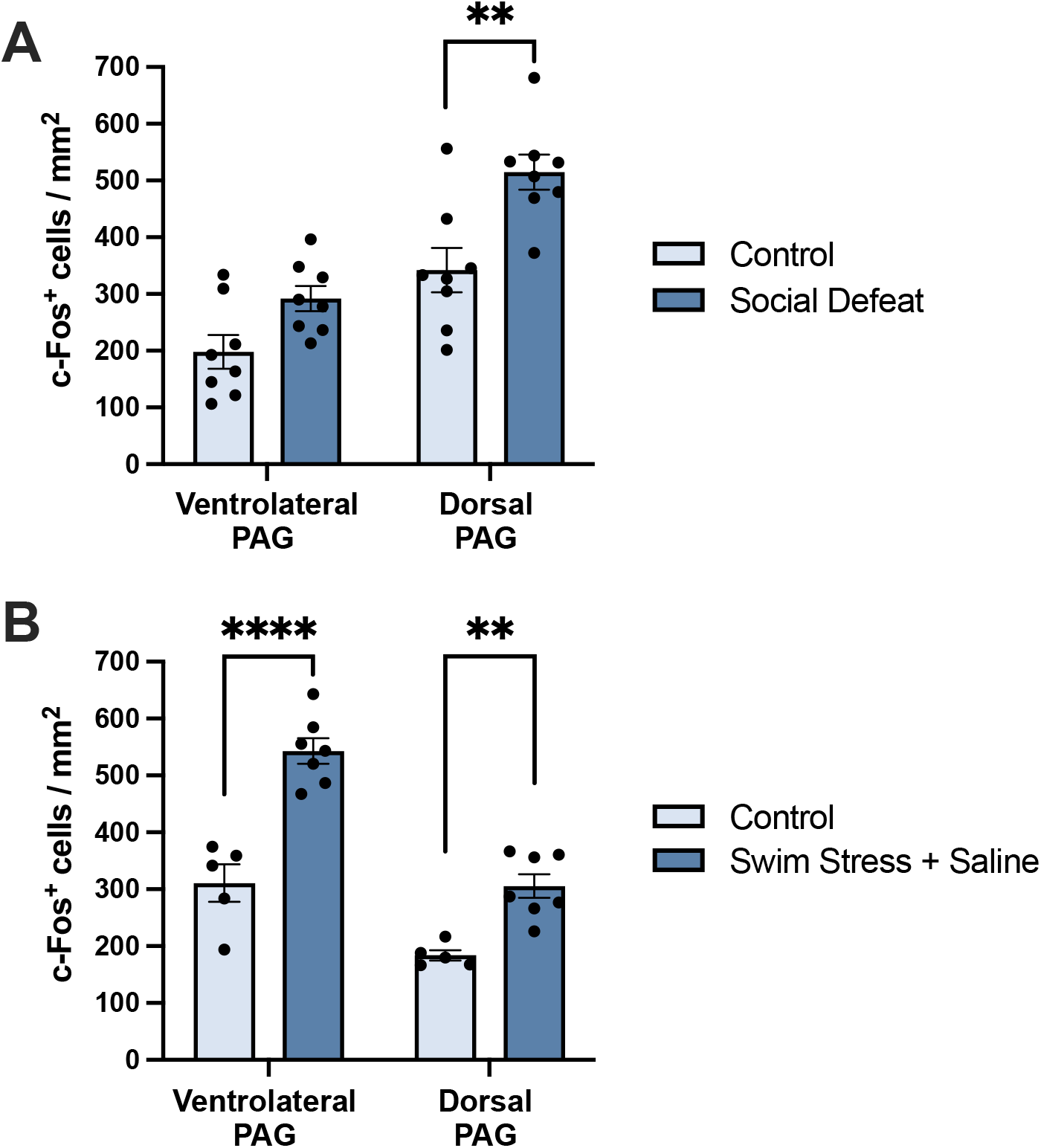
Effect of acute social defeat and swim stress on c-Fos immunoreactive neurons in the PAG. A) C-Fos immunoreactive cells in the PAG following social defeat (n=7) versus non-stressed controls (n=8). (B) C-Fos immunoreactive cells in the PAG following swim stress (n=7), swim stress with fluoxetine (n=7) versus non-stressed controls (n=6). Columns represent mean ± SEM values, with individual values indicated by closed circles. **** p<0.0001 *** p<0.001 ** p<0.01 * p<0.05.

### Mice with VGLUT3-deficient 5-HT neurons showed increased climbing during swim stress

Finally, we tested the causal role of glutamate co-released from 5-HT neurons in stress coping behaviour using genetically modified mice with VGLUT3 deletion targeted to 5-HT neurons (VGLUT3 cKO^5-HT^; ^25^). Specifically, we investigated the response of VGLUT3 cKO^5-HT^ mice to swim stress using climbing as a measure of active coping behaviour^40–42^. Previous studies have shown this behaviour to be increased by SSRI treatment^41,43^.

Firstly, we confirmed a loss of VGLUT3 in the DRN of VGLUT3 cKO^5-HT^ mice. Initial qPCR analysis demonstrated a 33.9 ± 5.7 % reduction of VGLUT3 mRNA in the midbrain raphe region of VGLUT3 cKO^5-HT^ mice compared to wildtype controls (t_(14)_=3.734, p=0.002; ***Fig 6B***). This effect was selective in that the VGLUT3 cKO^5-HT^ mice did not show altered expression of the vesicular monoamine transporter 2 (VMAT2) (t_(14)_=0.366, p=0.720; ***Fig 6B***). Then, immunocytochemistry confirmed a selective loss of VGLUT3 expression in DRN 5-HT neurons. Specifically, the number of TPH2/VGLUT3 co-labelled neurons in the ventral DRN of VGLUT3 cKO^5-HT^ mice was reduced by 62.6 ± 4.8 % compared to wildtype controls (t_(13)_=7.879, p<0.0001; ***Fig. 6C***). The TPH2 immunoreactive neuron count in the ventral DRN was not different between VGLUT3 cKO^5-HT^ mice and wildtype controls (t_(13)_=1.365, p=0.195; ***Fig 6C***), suggesting that the genetic deletion did not impact on the total number of 5-HT neurons.

**Figure 6.**
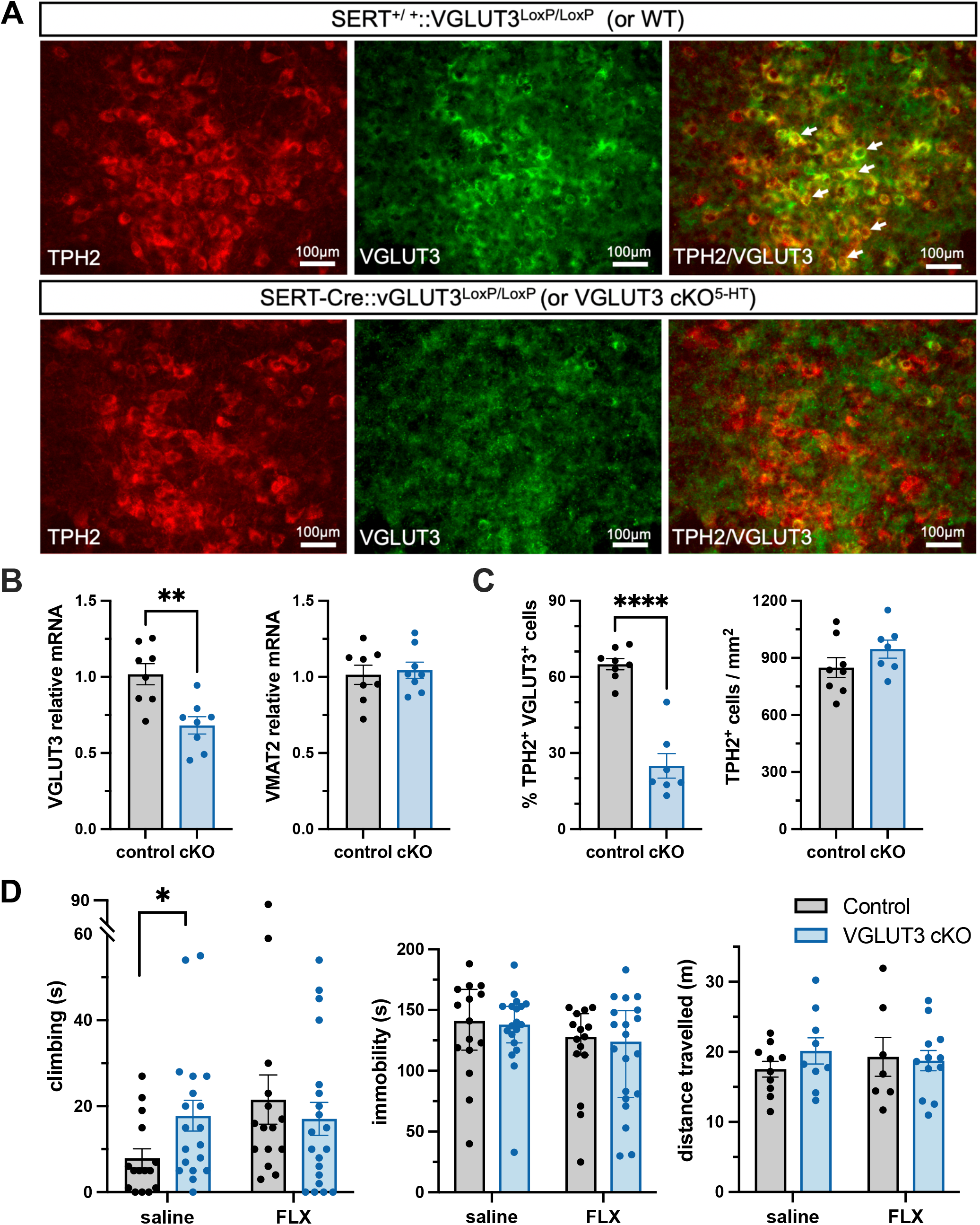
VGLUT3 cKO^5-HT^ mice; molecular characterisation and behavioural response to swim stress. (A) Representative image of TPH2/VGLUT3 double-labelled neurons (white arrows) in the ventral DRN of control mice (top) and VGLUT3 cKO^5-HT^ (bottom). (B) VGLUT3 and VMAT2 mRNA in the midbrain raphe region of VGLUT3 cKO^5-HT^ mice and littermate controls. (C) Number of TPH2/VGLUT3 co-labelled neurons (left) and TPH2 neurons (right) in the ventral DRN of VGLUT3 cKO^5-HT^ mice and littermate controls. (D) Performance of VGLUT3 cKO^5-HT^ mice (n=19-20) and littermate controls (n=15) during swim stress exposure. Columns are mean ± SEM values, except for climbing and immobility data that are medians with interquartile ranges. Individual values are indicated by closed circles. **** p<0.0001 ** p<0.01 * p<0.05.

The incomplete depletion of VGLUT3 may reflect cross-reactivity of our antibody with non-functional VGLUT3 protein fragments that may be transcribed following the conditional knockout. Also, even though the distribution of immunolabelling with this antibody closely matched that of VGLUT3 mRNA reported in previous in situ hybridisation studies^27^, we cannot exclude the possibility of a low level of non-specific labelling.

Prior to the behavioural testing of VGLUT3 cKO^5-HT^ mice we first established that pre-treatment of wildtype mice with fluoxetine increased time spent climbing when exposed to swim stress (Mann-Whitney U=6, p=0.016; ***Suppl. Fig. 1***) as observed previously^41,43^. Perhaps surprisingly, fluoxetine had no effect on time spent immobile (Mann-Whitney U=15, p=0.259; ***Suppl. Fig. 1***), but this has also been observed previously^42,44^. Although antidepressants normally reduce immobility in this paradigm, the C57BL/6 strain used here is generally less sensitive in this regard^45,46^ Moreover, the current small swimming chamber dimensions are reported to make it difficult to detect changes in immobility behaviour^47,48^.

Interestingly, in parallel with the effects of fluoxetine, when exposed to swim stress VGLUT3 cKO^5-HT^ mice also spent more time climbing versus littermate controls (Mann-Whitney U=77, p=0.042; ***Fig. 6D***) without having altered immobility time (Mann-Whitney U=131.5, p=0.917; ***Fig 6D***). Fluoxetine did not add further to the increase in climbing in the VGLUT3 cKO^5-HT^ mice, potentially because of a ceiling effect. The increase in climbing behaviour in the VGLUT3 cKO^5-HT^ mice was not associated with increased locomotor activity in that these mice showed similar levels of locomotion to their littermate controls in a separate locomotor test (effect of genotype: F_(1, 34)_=0.344, p=0.561; interaction: F_(1, 34)_=0.800, p=0.378; ***Fig 6D***).

The increase in climbing behaviour exhibited by VGLUT3 cKO^5-HT^ mice is evidence of enhanced escape-driven active coping behaviour which typically characterises the initial response to swim stress exposure^40^. Given our above immunocytochemical evidence that swim stress activates 5-HT-glutamate co-releasing neurons, it seems as if a deficiency in co-released glutamate in VGLUT3 cKO^5-HT^ mice promotes active coping behaviour. The predicted lack of co-released glutamate in the VGLUT3 cKO^5-HT^ mice would theoretically shift the 5-HT/glutamate balance at the synapse in favour of 5-HT. Interestingly fluoxetine, which also increased climbing behaviour, would also shift the 5-HT/glutamate balance in favour of 5-HT by selectively inhibiting 5-HT reuptake^49^. In other words, a switch in 5-HT/glutamate balance in favour of 5-HT may promote active stress coping behaviour.

The latter idea is consistent with a recent report that chemogenetic activation of ventral DRN-prefrontal cortex projecting 5-HT neurons increased active coping in mice exposed to swim stress^22^. Although the latter manipulation might be expected to release both 5-HT and glutamate, electrophysiological evidence from optogenetic studies^18^ suggests that 5-HT-glutamate co-release is frequency-dependent. This, glutamate was preferentially released at lower frequencies (1-2 Hz) and 5-HT was preferentially released at the higher frequencies (10-20 Hz). Therefore, chemogenetic activation may have preferentially released 5-HT resulting in increased active coping. Conversely, conditional TPH2 knockout from the same ventral DRN 5-HT neurons was found to increase immobility, supporting the hypothesis of the requirement for 5-HT in stress coping. Taken together, the evidence suggests that altered balance of 5-HT/glutamate in favour of 5-HT (i.e. away from glutamate and towards 5-HT-signalling pathways) may increase active coping and might therefore play a critical role in the behavioural response to stress.

A caveat of this hypothesis is the current lack of consensus regarding the mechanisms by which glutamate is co-released from 5-HT synapses^50^. The frequency-dependent nature of co-released glutamate and 5-HT evident in optogenetic studies^18^ indicates that 5-HT and glutamate are released from different vesicular pools. On the other hand, co-release from the same vesicular pools has also been suggested based on synergism between VGLUT3 and VMAT2^50^. In the latter scenario, VGLUT3 would promote vesicular loading of 5-HT^51^ in which case a reduction of VGLUT3 expression may decrease the vesicular content of both glutamate and 5-HT. Although this suggests that a loss of VGLUT3 in the VGLUT3 cKO^5-HT^ mice might disrupt the balance of glutamate-5-HT co-release less than expected, it is difficult to reconcile an increase in stress coping with an overall decrease in release of 5-HT in these animals (e.g.^22^).

The theory that a shift in balance of 5-HT/glutamate in favour of 5-HT increases coping would have implications in situations where this balance is altered, for example by environmental or genetic factors affecting the expression of VGLUT3 (but also VMAT2 or SERT). Interestingly, evidence suggests that the level of 5-HT-glutamate co-release may not be fixed but rather is plastic. For instance, changes in VGLUT3 expression in 5-HT neurons have been reported in rats exposed to chronic stress^52^ as well as during acquisition of generalised fear following acute stress^53^. More generally, VGLUT3 expression is reported to vary during neurodevelopment and early post-natal life^54,55^, and point mutations of the gene encoding VGLUT3 (Slc17a8) may result in a life-long alteration in VGLUT3 expression^56^. If the latter changes in VGLUT3 expression occur in 5-HT neurons and affect the balance of 5-HT-glutamate at the synapse, the present data suggest that they could impact on coping strategies and susceptibility to stress.

## Materials & Methods

### Animals

Mice were group housed (2-6 per cage) with littermates in individually ventilated cages, in a temperature-controlled room (21°C) with a 12 h light/dark cycle. Mice had *ad libitum* access to food and water, and cages were lined with sawdust bedding and contained cage enrichment (sizzle nests and a cardboard tube). Experiments were conducted during the light phase. Both female and male mice were used, except for the social defeat experiment which necessarily involved only males. Before each experiment mice were habituated to handling using a cardboard tunnel to minimise background stress^57^.

Most experiments utilised either C57BL/6J (Charles River, age 8-10 weeks) or transgenic mice with conditional VGLUT3 deletion targeted to 5-HT neurons (SERT-Cre::vGLUT3^LoxP/LoxP^, C57BL/*6*J background, age 8-17 weeks). The transgenic mice were generated by crossing VGLUT3^loxP/loxP^ mice (carrying a floxed allele of the exon 2 of Slc17a8) with a serotonin transporter (SERT)-Cre line^25^. SERT-Cre::VGLUT3^LoxP/LoxP^ were compared to control littermates (SERT^+/ +^::VGLUT3^LoxP/LoxP^ or WT). Retired male breeder CD1 mice (Charles River, age 22-30 weeks) were employed as resident aggressor mice for the social defeat experiments.

Experiments followed the principles of the ARRIVE guidelines and were conducted according to the UK Animals (Scientific Procedures) Act of 1986 with appropriate personal and project licence coverage.

### Swim stress paradigm

Mice were randomly allocated to 1 of 3 experimental groups by stratified randomisation: i) saline; ii) saline + swim stress; and iii) fluoxetine (10 mg/kg) + swim stress. Mice were removed from their home cages and single-housed in a clean cage before and after undergoing a single exposure to swim stress. Saline or fluoxetine were injected i.p. 30 min prior to a swim stress.

During the last 5 min prior to swim stress, mice were placed in a clean but familiar cage, and their locomotor activity was recorded via an overhead camera for offline tracking using ANY-maze (Stoelting Europe) tracking software.

For swim stress, mice were placed individually for 6 min in a glass cylinder (height 25 cm, diameter 12 cm) containing water (height 20 cm) maintained at 20°C, as described previously^58,59^. A video camera was mounted in front of the cylinder and recordings were used for offline manual scoring by an experimenter blind to treatment. Climbing and immobility were timed during the final 4 min of stress exposure. Climbing was defined as placement of the front paws on the glass surface above water level^40,41^, whilst immobility was rated as the absence of escape-oriented behaviours. After the test, animals were towel-dried and placed in a heated cage until fully dry.

Ninety min after swim stress mice were deeply anesthetised prior to perfusion and collection of brain tissue for c-Fos immunohistochemistry (***Fig. 7***). This timescale was chosen to allow for optimum c-Fos expression before tissue collection^60^.

**Figure 7.**
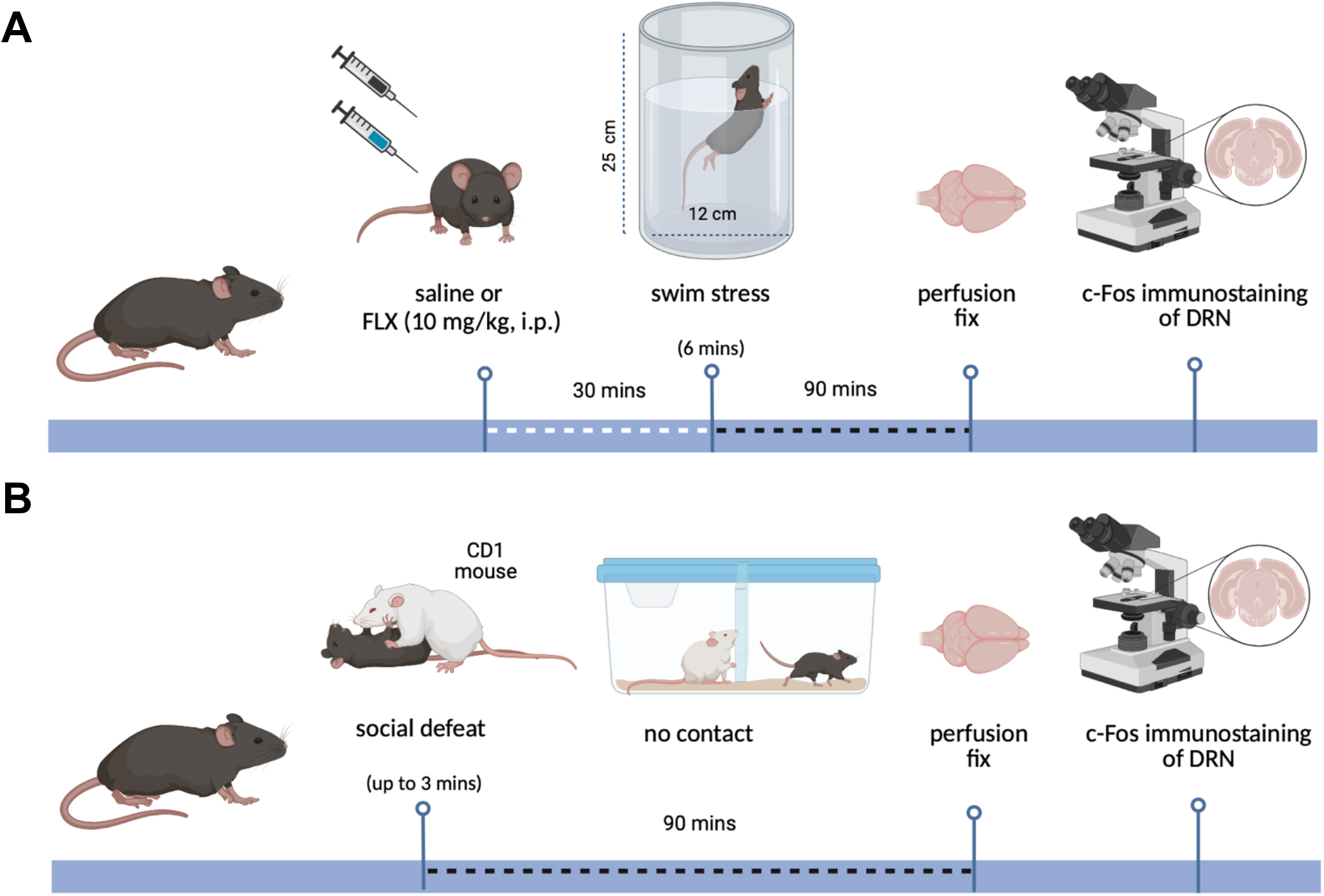
Experimental timeline. (A) Timeline of swim stress (top) and social defeat (below) experiments. Abbreviations: fluoxetine (FLX), dorsal raphe nucleus (DRN). Created with BioRender.com.

### Social defeat paradigm

Male mice (C57BL/6J) were randomly allocated to two experimental groups by stratified randomisation: i) control and ii) social defeat. On the day of social defeat, mice were removed from their home cage and single-housed in a clean but familiar cage. Control mice remained in the clean cage for 90 min^61^. In the ‘social defeat’ condition, an intruder mouse was placed in the home cage of a territorially-dominant, aggressive resident mouse and subject to brief social defeat (as defined below). The intruder was then separated from the resident by a perforated acrylic partition which allowed auditory, visual, and olfactory interaction with the resident but no physical contact^61^. After 90 min mice were deeply anaesthetised and perfused (see below). The resident-intruder interaction was recorded with an overhead camera for offline behavioural analysis using ANY-maze software (Stoelting Europe).

### Resident mouse training and selection

Resident mice were selected based on a persistent level of aggression as previously described^61^. Briefly, on three consecutive days an intruder mouse was placed in the cage of a resident mouse for up to 3 min or until the latter was ‘socially defeated’. Social defeat was defined as a clear pin-down and/or a supine posture of the intruder. Each resident mouse interacted with a different intruder mouse daily. All interactions were filmed and video analysis of the latency to attack and number of attacks allowed selection of resident mice that consistently attacked within the first 20 s of the resident-intruder interaction.

### Immunohistochemistry and microscopy

Mice were deeply anesthetised by i.p. injection of with sodium pentobarbital (90 mg/kg; Euthatal) and intracardially perfused with 4 % paraformaldehyde in phosphate-buffered saline (PBS). Brains were then dissected, post-fixed by immersion in the same fixative for 48 h, cryoprotected in PBS containing 30 % sucrose and frozen at −80°C until sectioning.

Cryostat-cut coronal brain sections (30 μm; Bright LOFT cryostat) were taken at the level of the DRN (Bregma: −4.6^26^ and stored in antifreeze (30 % glycerol, 30 % ethylene glycol, in PBS) at −20 °C prior to processing for immunohistochemistry as previously described^62^. In brief, sections were incubated overnight with the following primary antibodies: rabbit anti-c-Fos (1:1000, Abcam), goat anti-TPH2 (1:1000, Abcam), guinea pig anti-vGLUT3 (1:500 dilution, Synaptic Systems). The secondary antibodies used for protein visualization were the following: rabbit AF488 (1:1000, Invitrogen), guinea pig Cy3 (1:1000, Jackson Immune Research), goat AF647 (1:1000, Abcam). Cell nuclei were stained using DAPI (1:1000, 5 min).

Images were visualised using an epi-fluorescent microscope (Olympus BMAX BX40) and acquired with ImageJ Micromanager v1.4 (500 ms exposure). Sections were imaged at 20x magnification for the ventral DRN, dorsal DRN and MRN, and at 10x for the entire DRN, lateral wings, ventrolateral and dorsal PAG^26^. Cell counting and quantification of colocalization was performed by an experimenter blind to treatment employing the ImageJ Software package.

For each mouse, the mean cell count of 3 sections was used for statistical analysis. C-Fos immunoreactive cells with colocalised with DAPI immunoreactivity were defined as neurons. Colocalization of DAPI and TPH2 immunoreactivity identified 5-HT neurons, whist colocalization of TPH2 and VGLUT3 identified 5-HT-glutamate co-releasing neurons.

### Drugs

Fluoxetine hydrochloride (Stratech A2436-APE) was dissolved in 0.9 % sodium chloride in at 2 mg/ml and administered i.p. at a dose of 10 mg/kg. Control mice received saline in a volume of 2 ml/kg. All solutions were prepared fresh daily. Fluoxetine dose and administration protocol were based on previous studies^45,46^.

### qPCR analysis

For PCR analysis, the midbrain raphe region was dissected from frozen tissue sections (1 mm). RNA was extracted (Qiagen RNeasy Mini Kit) using the TRIzol method^63^ and eluted into 20 μl of RNase-free water. DNA conversion and qPCR were conducted as described previously^64^. In brief, conversion to cDNA was achieved using a high-capacity cDNA Reverse Transcription Kit (Life Technologies) and T100 Thermocycler (Bio-Rad). QPCR was performed (800 ng RNA) using a LightCycler® 480 instrument (Roche Diagnostics) with the following primers (300 nM): VGLUT3 (specifically targeting the exon 2) 5’-CGATGGGACCAATGAAGAGGA-3’ and 5’-CAGTCACAGACAGGGCGATG-3; VMAT2 5’-CATCACGCAGACTTGAAAGAC-3’ and 5’-CGCCTCGCCTTGCTTATCC-3’^65^. GAPDH was used as the reference gene (Santa Cruz Biotechnology). Reactions (384 well-plates, 10 μl reaction volume, 5 μl Precision®PLUS qPCR Master Mix with SYBRgreen, 25 ng cDNA) used the following cycle: enzyme activation for 2 min at 95 °C, 40 cycles of 10 s at 95 °C, 1 min at 60 °C, then held at 4 °C. Samples were run in triplicate and 2^-ΔΔCT^ was calculated for each sample, where ΔCT = CT_target gene_ – CT_reference gene_. Data were analysed as fold-change in gene expression relative to the control group.

### Statistical Analysis

The Shapiro-Wilk test for normality was applied to all data sets. If data were normally distributed, then a t-test, one-way or two-way ANOVA were used followed by Tukey’s or Šídák’s post hoc tests as appropriate. Specifically, when c-Fos data was analysed across multiple regions, repeated-measures two-way ANOVA was employed for balanced data whereas a repeated-measure mixed-effect model was used for datasets with missing values. If the data were non-parametric then a single or multiple Mann-Whitney test was employed, with Holm-Šídák correction for multiple comparison. GraphPad Prism was used for all analysis and plotting of graphs. Parametric data are presented as mean ± standard error of the mean (SEM) values, whilst non-parametric data are presented as median ± interquartile range values; p<0.05 was considered statistically significant.

## Abbreviations

5-HT: 5-hydroxytryptamine
DRN: dorsal raphe nucleus
FLX: fluoxetine
MRN: median raphe nucleus
PAG: periaqueductal gray
SSRI: serotonin reuptake inhibitor
TPH2: tryptophan hydroxylase 2
VGLUT3: vesicular glutamate transporter 3
VMAT2: vesicular monoamine transporter 2

## Author information

SG performed experiments, analysed the data, and contributed to writing the manuscript. CF and PD performed *ex vivo* tissue analysis. HC contributed to behavioural experiments. SEM contributed to manuscript preparation. TS contributed to the conception and design of the work, drafting, and revising the manuscript, and interpretation of data.

## Acknowledgments

SG was supported by a studentship from the Oxford-MRC Doctoral Training Programme. HC was supported by a Wellcome Trust PhD Studentship in Basic Science (HC; grant no. 219982/Z/19/Z).

