## Supplementary information for "Evidence for a role of 5-HT-glutamate co-releasing neurons in acute stress mechanisms"

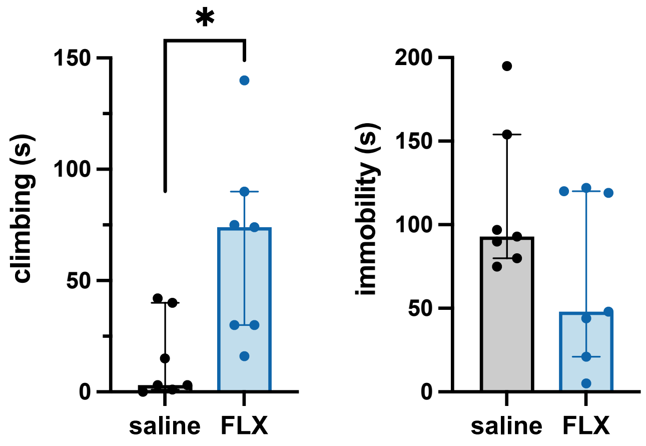


**Supplementary Figure 1 | Behaviour of mice exposed to swim stress with and without fluoxetine.** Time spent climbing (left) and immobile (right) during swim stress exposure. Columns represent median ± interquartile range values, with individual values indicated by closed circles. Groups were saline (n=7) and fluoxetine (FLX, n=7). * $p<0.05$.

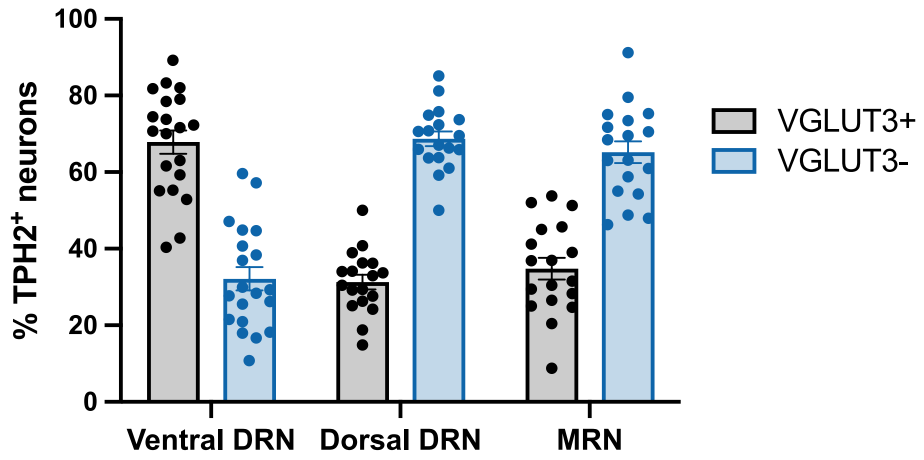


**Supplementary Figure 2 | Colocalization of TPH2 and VGLUT3 in neurons of mouse raphe regions.** Percentage of TPH2 neurons that co-localised with VGLUT3, or were

VGLUT3 immunonegative (n=17-20). Bars represent mean ± SEM values, with individual values are indicated by closed circles.

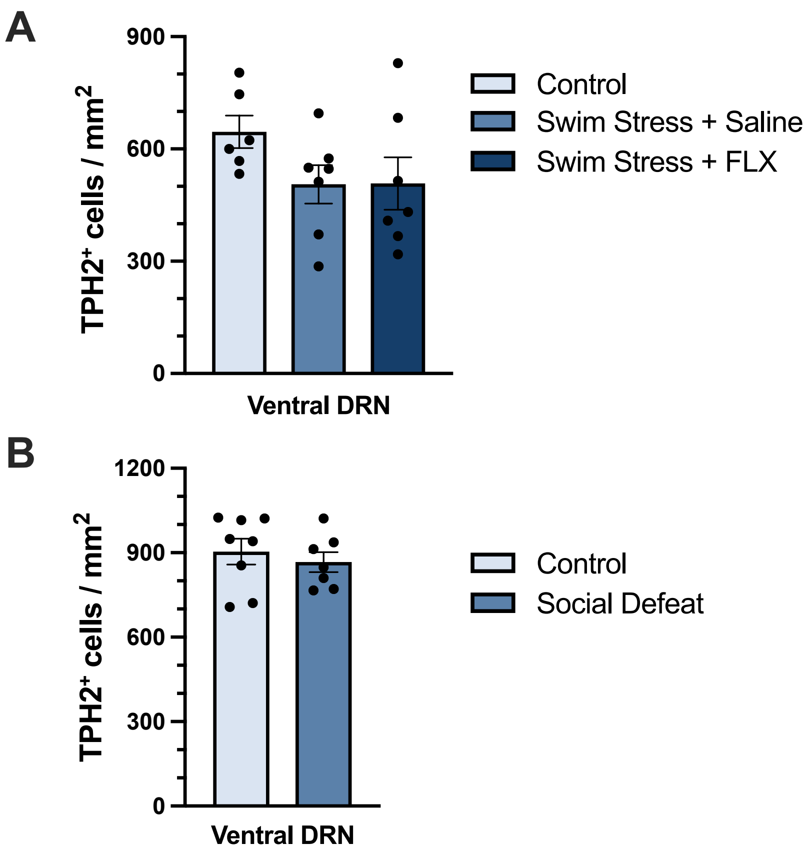


**Supplementary Figure 3 | TPH2 expression in ventral DRN of mice exposed to swim stress and social defeat.** A) Number of TPH2 immunoreactive neurons following swim stress with or without fluoxetine; groups were control (n=6), saline + swim stress (n=7) and 10mg/kg fluoxetine + swim stress (n=7). (B) Number of TPH2 immunoreactive neurons following a single episode of social defeat; groups were control (n=8) and social defeat (n=7). Bars represent the mean ± SEM values with individual values are indicated by closed circles.
